# The Alzheimer’s Disease-associated gene CD2AP regulates permeability of endothelial cells and transendothelial migration of immune cells

**DOI:** 10.64898/2026.10.01.755982

**Authors:** Michael D Onken, John A Cooper

**Affiliations:** Department of Biochemistry & Molecular Biophysics, Washington University School of Medicine, St. Louis, MO 63110, USA

**Keywords:** CD2AP, Transendothelial migration, Endothelial cells, NK cells

## Abstract

Variants at the CD2AP locus, which encodes a branched-actin regulator, are associated with late-onset Alzheimer’s disease (AD), and CD2AP-deficient mice show increased blood-brain barrier (BBB) permeability. CD2AP is enriched in endothelial cells, but its function there is poorly understood. In endothelium, branched actin networks maintain cell-cell junctions and are also required for immune cells to cross through individual endothelial cells (transcellular transendothelial migration, TEM). We previously showed that the Arp2/3 activators N-WASP and WAVE2 contribute to these processes, but the capping protein (CP)-interacting proteins involved are unknown. Here we tested whether CD2AP, a CP-interacting protein recruited to cell-cell contacts, regulates both junction integrity and TEM. In primary human microvascular endothelial cells, shRNA knockdown of CD2AP increased TNFα-induced monolayer permeability, measured by transendothelial electrical resistance, and altered VE-cadherin junction morphology. Unexpectedly, on soft hydrogels the same knockdown reduced TEM of NK-92 MI cells, lowering the proportion of transcellular events. Thus, loss of CD2AP weakens the endothelial barrier yet impairs a distinct, actin-dependent route of immune-cell passage. These findings suggest that CD2AP risk variants may affect the BBB through more than permeability, and they identify endothelial immune-cell trafficking as a testable link between CD2AP and AD.

## INTRODUCTION

Alzheimer’s Disease (AD) is the most prevalent cause of late-onset dementia, characterized by the accumulation of amyloid-beta (Aβ) protein into extracellular plaques produced by neurons and oligodendrocytes (Kamenetz, 2003; Rajani, 2024). In response, astrocytes and microglia become activated to clear these plaques and shield neurons from further damage – though this protective response can gradually give way to chronic neuroinflammation (De Strooper, 2016). While much of the translational and pre-clinical research has centered on reducing amyloid plaque formation, the roles of microglial activation in AD progression have been recognized for some time (Sarlus, 2017). Likewise, microglial cells regulate permeability of the blood-brain barrier (BBB) during inflammation, which plays a key role in AD progression: crosstalk between activated microglia and the BBB endothelium changes endothelial permeability to allow small molecules to cross and to facilitate the extravasation of immune cells that contribute to the inflammatory process (Thurgur, 2019). Thus, a properly functioning BBB is essential to proper microglial function and homeostasis and to slowing the progression of AD.

Over a decade ago, two large companion Genome-Wide Association Studies identified the *CD2AP* gene as a strong genetic predictor for AD (Hollingworth, 2011; Naj, 2011). Early investigations into this link focused on neuronal Aβ protein production and deposition, but CD2AP knockdown in neurons had only subtle effects on Aβ levels and no effects on plaque formation in mice (Liao, 2015). Instead, analyses of RNAseq datasets show low-level expression of CD2AP in neurons (Lein, 2007) but 5-fold higher expression in endothelial cells (Zhang, 2014), and further immune-histochemical studies showed that the strongest CD2AP staining in the brain was on endothelial cells in the brain vasculature (Li, 2000; Uhlen, 2010; Lehtonen, 2008). Moreover, CD2AP knockout mice showed a significant increase in BBB permeability to small molecules suggesting a cerebrovascular role for CD2AP in Alzheimer’s disease risk (Cochran, 2015).

In endothelial cells, branched actin networks maintain cell-cell junction morphology, dynamics, and permeability and are required for transcellular trans-endothelial migration (TEM) of immune cells (Onken, 2014). Formation of functional branched actin networks requires the Arp2/3 complex to generate branchpoints and Capping Protein (CP) to reduce actin filament elongation in favor of branching (reviewed in Hammer, 2026). The Arp2/3 complex is activated by membrane-associated nucleation promoting factors, and in endothelial cells, activation of Arp2/3 by N-WASP is required for junction integrity, whereas activation by WAVE2 is involved in TEM (Mooren, 2014; Mooren, 2015). Likewise, CP is activated by membrane-associated proteins containing CP-interacting (CPI) domains, but in contrast to Arp2/3, the specific CPI-containing proteins required for junction integrity TEM in endothelial cells are not fully established. CD2AP is an SH3-adapter protein with a CPI domain that is recruited to the plasma membrane at sites of cell-cell contact (van Duijn, 2010) where it stabilizes cell-cell junctions by activating local branched actin networks (Tang, 2013) through its CPI domain (Bruck, 2006; Edward, 2015). The main goal of this study was to establish CD2AP as a vital regulator of actin networks in endothelial cells, specifically at junctions between adjacent endothelial cells responsible for vascular permeability and at contact sites with immune cells required for TEM.

## RESULTS

### CD2AP knockdown impairs junction maturation in TNFα-treated endothelial monolayers

To test the role of CD2AP in endothelial cells, we reduced its expression in primary human microvascular endothelial cells (HMVECs) using lentiviral shRNA. Two independent constructs (#1 and #4) each reduced CD2AP protein by approximately 80% relative to a scrambled control (Fig. 1A), and each were used throughout this study. We grew HMVECs on soft polyacrylamide hydrogels, where they form mature monolayers within three days (Onken, 2014), and treated them overnight with vehicle or TNFα to model inflammation. Fixed monolayers were stained for VE-cadherin, a marker of adherens junctions, and for F-actin.

**Figure 1.**
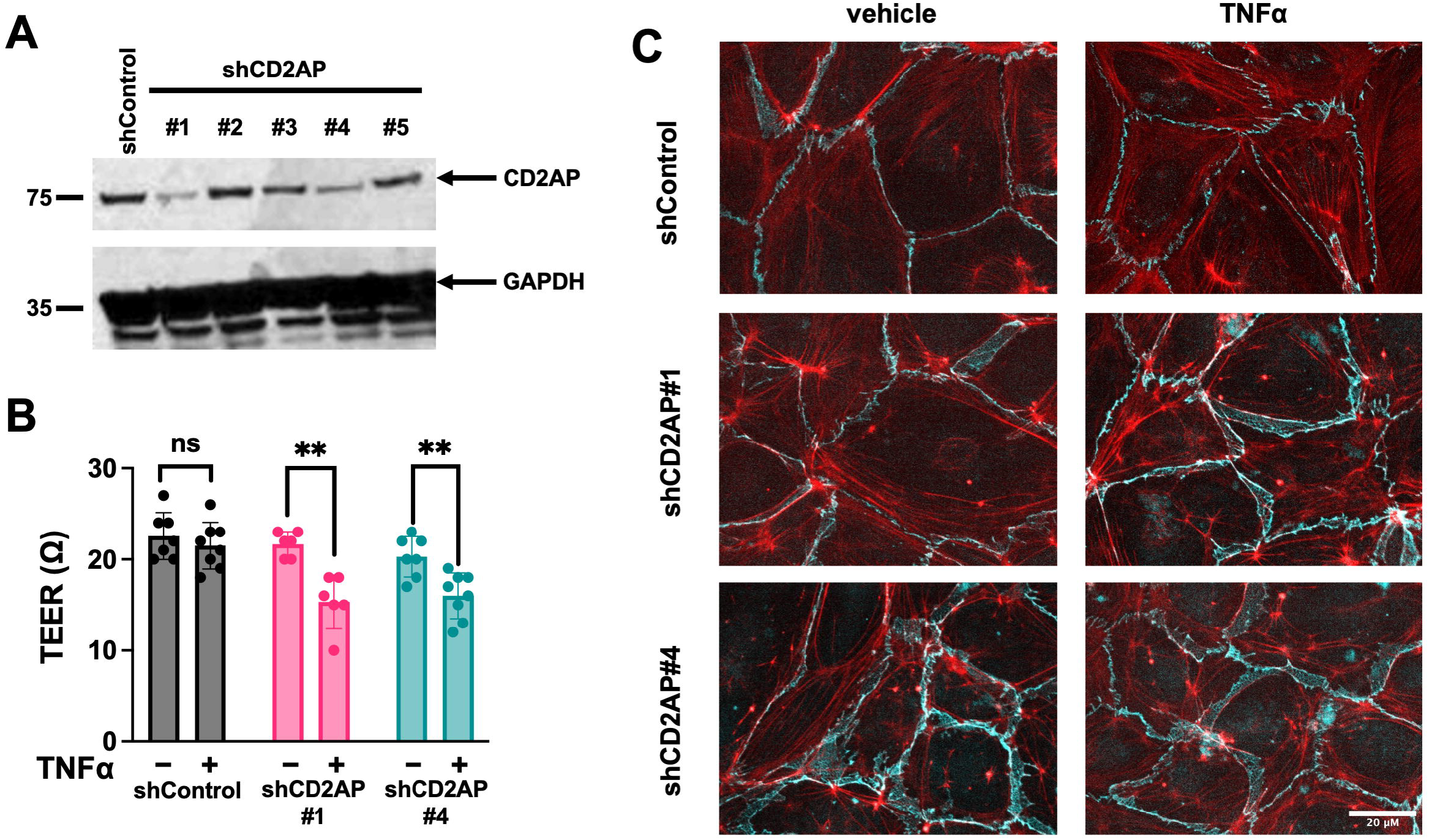
CD2AP knockdown alters permeability of human microvascular endothelium. **A)** Immunoblot for CD2AP in HMVECs stably expressing control or CD2AP short-hairpin RNA interference constructs. 5 different short-hairpins were screened and two, #1 and #4, showed significant knockdown of endogenous CD2AP expression. Immunoblot for GAPDH was performed on the same membrane as a loading control. **B)** Trans-endothelial electrical resistance (TEER) was used to measure permeability of HMVEC stables grown on 0.4 µm pore filters in transwells. Measurements were made after overnight treatment of monolayers with TNFα or vehicle. Data were collected from 3 independent experiments, each starting from a different batch of uninfected HMVECs (* p < 0.01, Welch’s t-test). **C)** Immunofluorescence histology was performed on HMVEC monolayers stably expressing short-hairpin constructs and treated overnight with TNFα or vehicle. Anti-VE-cadherin (Cyan) and phalloidin (F-actin; Red) were used to identify cell junctions. Scale bar is 20 µm.

In fixed monolayers, mature endothelial junctions appear as narrow, straight lines of VE-cadherin, whereas immature, remodeling junctions are wider and less linear (Huveneers, 2013; Onken, 2014; Kim, 2018). After TNFα treatment, CD2AP knockdown produced wider, less linear VE-cadherin junctions than control (Fig. 1B). This is consistent with impaired junction maturation in the absence of CD2AP.

### CD2AP knockdown increases TNFα-induced permeability

Because junction maturation is a determinant of barrier function, we next asked whether CD2AP knockdown affects monolayer permeability, measured as transendothelial electrical resistance (TEER). After overnight TNFα treatment, CD2AP-knockdown monolayers had significantly lower resistance than control monolayers (Fig. 1C). TEER differences between vehicle-treated monolayers were not significant. This agrees with the increased BBB permeability seen in CD2AP knockout mice (Cochran, 2015) and shows that loss of CD2AP in endothelial cells alone is sufficient to weaken the barrier under inflammatory conditions.

Together, these results indicate that CD2AP helps endothelial junctions mature and resist inflammatory disruption. Junction integrity is one of two actin-dependent endothelial functions we have studied (Mooren, 2014; Mooren, 2015), the other being support of immune-cell transmigration. We therefore asked whether CD2AP also contributes to TEM.

### CD2AP knockdown reduces immune-cell transendothelial migration

Given the role of CD2AP at endothelial junctions, we expected its loss to make the monolayer easier for immune cells to cross. To test this, we measured TEM of NK-92 MI natural killer cells across TNFα-treated HMVEC monolayers on soft hydrogels, using time-lapse DIC microscopy. NK-92 MI cells are a well-characterized system in our hands for distinguishing the routes of TEM (Mooren, 2014; Mooren, 2015) (Fig. 2A). NK-92 MI cells were added at time zero and imaged every 20 seconds, and a transmigration event was scored when a rounded NK-92 MI cell was seen passing through the monolayer and spreading beneath it, between the endothelium and the hydrogel (Fig. 2B-C). Contrary to that expectation, CD2AP knockdown reduced total TEM relative to control (Fig. 3A-B) by 32% for shCD2AP#1 and 46% for shCD2AP#4 (p<0.01, Welch’s t test, n = 4 independent experiments). Thus, loss of CD2AP has opposite effects on two endothelial functions: it weakens the barrier, but it also makes the monolayer harder for immune cells to cross.

**Figure 2.**
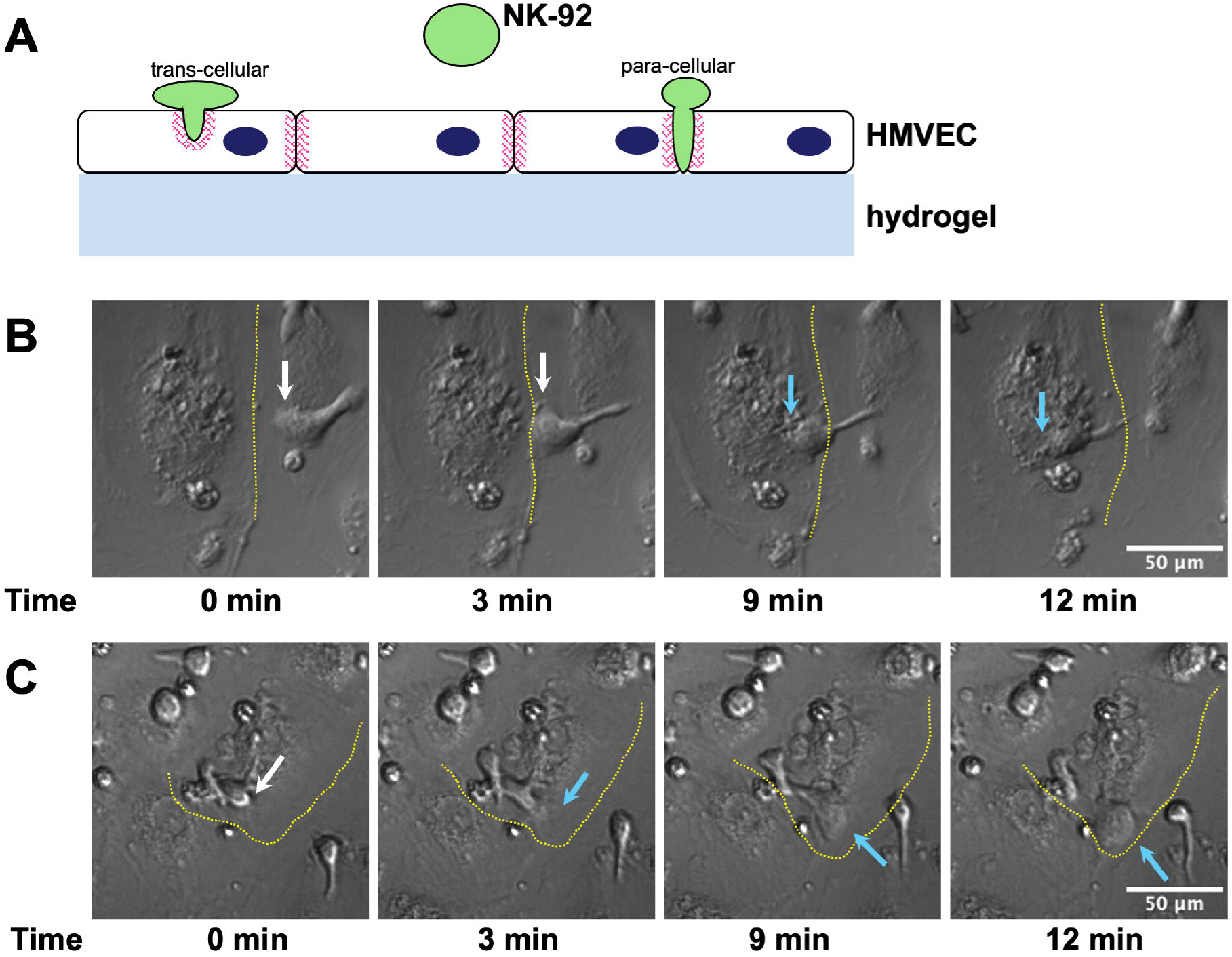
Model system for measurement of Trans-Endothelial Migration (TEM) of NK cells across HMVEC monolayers. **A)** Schematic representation of the model: HMVEC monolayers are formed on polyacrylamide soft substrates (hydrogel); NK-92 cells are added to the medium over the monolayers immediately before imaging; and transmigration is scored by time of initiation and whether transmigration occurs at a junction between two cells (para-cellular) or through an endothelial cell away from a junction (trans-cellular). **B)** Representative time points showing an NK-92 cell crawling on a TNFα-activated HMVEC monolayer. A dotted yellow line is superimposed to emphasize a junction between two HMVEC cells. The white arrow indicates the leading edge of the NK cell during crawling above the monolayer. The cyan arrow indicates the leading edge of the NK cell under an endothelial cell. Scale bar is 50 µm. **C)** Representative time points from a different experiment showing an NK-92 cell crawling on a TNFα-activated HMVEC monolayer. A dotted yellow line is superimposed to emphasize the junctional perimeter around an HMVEC. The white arrow indicates the leading edge of the NK cell during initial interaction with the HMVEC. The cyan arrow indicates the leading edge of the NK cell as it passes through and beneath the endothelial cell. Scale bar is 50 µm.

**Figure 3.**
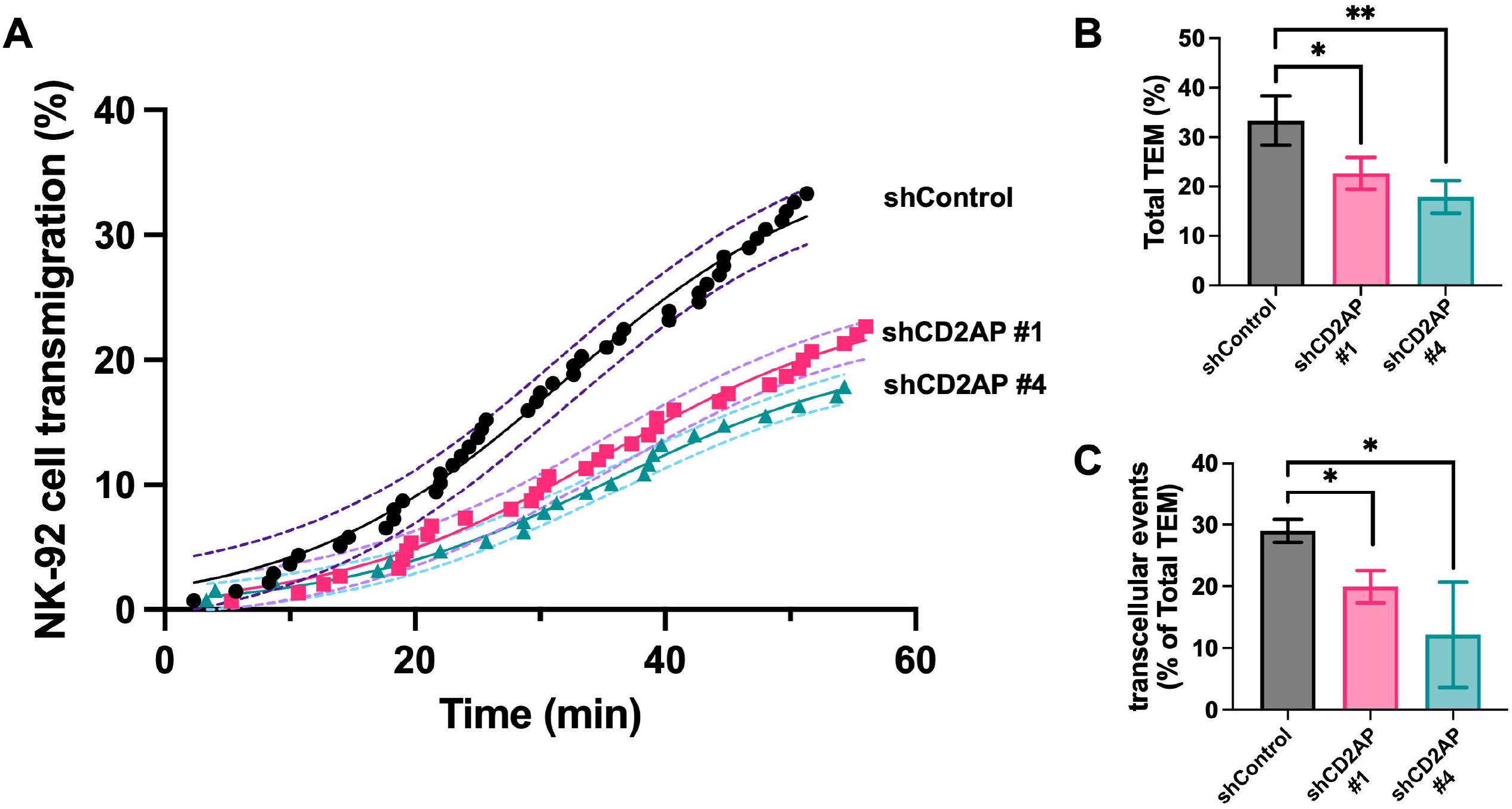
CD2AP knockdown reduces TEM of NK cells across HMVEC monolayers. **A)** Representative results of TEM by NK-92 cells on HMVEC monolayers. Graph shows combined results from 3 technical replicates of each condition from one experiment. Short-hairpin constructs are indicated for each trace. Time indicates time of initiation of transmigration for each point. **B)** Data collected from 3 independent experiments show significant reduction of TEM by NK-92 cells through monolayers with CD2AP stably knocked down compared to control monolayers (* p < 0.01; ** p < 0.001, Welch’s t-test, n = 4). **C)** Route of TEM was noted for each event in all movies in B) based on migration visibly away from cell junctions and calculated as a percentage of total TEM (* p < 0.01, Welch’s t-test, n = 4).

### CD2AP is required for the transcellular route of TEM

Immune cells can cross an endothelial monolayer between cells (paracellular route) (example in Fig. 2B) or through the body of an endothelial cell (transcellular route) (example in Fig.2C). Both require branched actin, but the transcellular route demands more extensive remodeling of the endothelial cell than opening and resealing a junction does (Onken, 2014). We therefore re-analyzed the same movies and classified each completed event by route. The percentage of events that were transcellular was significantly lower in CD2AP-knockdown monolayers than in control (Fig. 3C; p<0.01, Welch’s t test, n = 4 independent experiments). Because the barrier is leakier in these cells, the reduction in transcellular events is not explained by tighter junctions, which suggests that CD2AP contributes to transcellular TEM by a mechanism separate from its role in junction maturation.

This result differs from a previous study reporting that CD2AP loss increased transcellular relative to paracellular diapedesis (Schaefer, 2017). That study used HUVECs, large-vein endothelium grown on glass, whereas ours used microvascular cells on soft hydrogels. Both endothelial origin and substrate stiffness influence which route immune cells take (Onken, 2014), so the two findings need not conflict, but they do show that the role of CD2AP in TEM depends on context.

## DISCUSSION

### CD2AP acts at two sites, with different consequences

Our results show that in microvascular endothelial cells, CD2AP supports both the junction barrier and the transcellular passage of immune cells. One interpretation is that the same capping-protein–interacting adapter is used at two locations, at cell-cell junctions and at sites of immune-cell contact, to build the branched actin networks each requires. CD2AP binds ICAM-1 (Schaefer, 2017), which would place it at leukocyte contact sites, but our data do not show that CD2AP localizes there or that its actin-regulatory activity is needed. Testing whether the CP-interacting (CPI) domain is required for each phenotype, for example by rescuing with a CPI-mutant CD2AP, is the next step.

### Relevance to Alzheimer’s disease

CD2AP is an AD risk locus, is highly expressed in brain endothelium, and its deficiency increases BBB permeability in mice (Cochran, 2015). Our data agree with the permeability phenotype and show that it originates in endothelial cells. They also raise a new possibility: that CD2AP affects immune-cell trafficking across the BBB. Whether reduced trafficking would help or harm in AD is unknown. Reduced infiltration might limit neuroinflammation, or it might impair immune surveillance and clearance. Because the AD risk variant rs9349407 lowers CD2AP expression without altering the protein’s function (Hollingworth, 2011; Naj, 2011), partial knockdown is a more relevant model than complete knockout, and these experiments provide a basis for testing patient-derived or variant-carrying endothelial cells.

### Limitations

These experiments used dermal microvascular endothelial cells, not brain endothelium, and NK-92 MI cells as a surrogate for infiltrating immune cells, so whether the same phenotypes occur at the BBB and with other cell types, such as T cells and monocytes, remains to be shown. The 80% knockdown is also greater than the reduction expected from a risk allele, so an effect at more modest reductions should be tested.

## METHODS

### Cell Culture

Cells were cultured at 37°C in 5% CO_2_. Human dermal microvascular ECs (HMVECs) were obtained from Lonza (Allendale, NJ) and were cultured in EBM-2 base medium supplemented with the EGM-2 MV kit.

HMVECs were not used after 9 passages. Human NK-92 MI cells (ATCC, Manassas, VA) were cultured in alpha-MEM containing 12.5% Fetal Bovine Serum, 12.5% Horse Serum, 2 mM L-glutamine, 1.5 g/L sodium bicarbonate, 0.2 mM inositol, 0.02 mM folic acid, and 0.1 mM 2-mercaptoethanol. For transmigration assays and immunofluorescence studies, HMVECs were grown on polyacrylamide hydrogels, which were prepared as described (Onken, 2014). Briefly, 20-mm glass-bottom culture dishes were coated with 3-aminopropyltriethoxysilane and activated by treatment with 0.5% glutaraldehyde for 30 min. Polyacrylamide mixtures were prepared in PBS with 7% acrylamide, 3% acrylamidopropyltrimethyl ammonium chloride, and 0.2 % bis-acrylamide, and 20 μL of the acrylamide mixture was placed in the center of the treated glass-bottom dish. An 18-mm coverslip was placed on top to create a smooth, even gel, sandwiched between the coverslip and the glass-bottom dish. The acrylamide was allowed to polymerize for 30 min at RT, the coverslip was removed, and the gel was washed three times with PBS. The substrates were coated with fibronectin (10 µg/mL in PBS) by 30 min incubation at 37ºC and washed with PBS. 2 × 10^4 HMVECs were added to each dish in 250 µL of growth medium directly to the soft substrate and allowed to form monolayers overnight. 2 mL of growth medium was added the next morning, and monolayers were given at least two days to mature.

Tumor necrosis factor alpha (TNFα) was obtained from Gibco ThermoFisher (Gaithersburg, MD, USA) and fibronectin was obtained from Sigma-Aldrich (St Louis, MO, USA). Five shRNA lentiviral expression constructs targeting CD2AP and one shRNA scrambled control construct were purchase from the MISSION® shRNA collection of MilliporeSigma (St Louis, MO, USA) as lentiviral particles. The five CD2AP shRNA constructs were TRCN0000119097 (#1), TRCN0000119098 (#2), TRCN0000119099 (#3), TRCN0000119100 (#4), and TRCN0000119101 (#5). The MISSION® Non-Mammalian shRNA Control was SHC002.

### Immunoblots

Confluent HMVECs were washed with room temperature PBS and lysed by adding ice-cold 1% NP-40 lysis buffer into culture dishes and incubating on ice for at least 30 min. Cell lysates were harvested by scraping the dish, and they were centrifuged at 13,000 rpm for 15 min at 4°C. The supernatant was incubated with SDS-loading buffer, boiled for 5 min, electrophoresed on SDS-PAGE gels, and transferred to PVDF membrane (EMD Millipore Corp., MO, USA) at 0.22 A overnight. The membrane was incubated in blocking buffer (4% BSA in TBST buffer) and then in a primary antibody mixture overnight at 4°C. Primary antibodies were a polyclonal rabbit anti-CD2AP (Cell Signaling) and a monoclonal mouse anti-human GAPDH (clone 6C5) obtained from Abcam (Cambridge, MA, USA). After washing with PBS, the membrane was incubated in horseradish-peroxidase-conjugated secondary antibodies (Life Technologies) at room temperature for 1 hr. The membrane was incubated with enhanced chemiluminescence solution for 5 min at RT and developed for autoradiography.

### Immunostaining

Fixation was carried out by adding 2X fixative (PBS containing 4% paraformaldehyde and 0.2% glutaraldehyde) directly to live cells in tissue culture medium. After 15 min at 37ºC, cells were rinsed three times in PBS to remove all fixative and then permeabilized with 0.1% Triton X-100 in PBS for 3 min, washed with PBS, and blocked with 2% fish gel (teleostean gelatin; Sigma) in PBS. Primary and secondary antibodies were diluted in 2% fish gel. Primary antibody was rabbit anti-VE-cadherin (Cell Signaling, Danvers, MA). F-actin was visualized using Alexa-fluor conjugated phalloidin (Life Technologies). Secondary antibodies were Alexa-fluor conjugates (Life Technologies), and the mounting agent was ProLong Gold with DAPI (Molecular Probes ThermoFisher, Eugene, OR, USA). Cells were imaged with an inverted microscope (Olympus IX83) using 40X and 100X objectives. Images of monolayers were analyzed using FIJI (Schindelin, 2012).

### Transendothelial Electrical Resistance assays

HMVECs were seeded at 5×10^4 cells in 100 µL on the apical side of 0.4 µm porosity inserts (BD Falcon, Corning) that had been coated with 10 µg/mL Fibronectin for at least 1 h at 37°C prior to adding the cells. The cells were grown 3-4 days in EGM-2MV growth medium (Lonza). Trans-endothelial electrical resistance (TEER) of monolayers was measured using an EVOM-2 epithelial voltohmmeter with ENDOHM-6 transwell cup electrodes. Monolayers were considered complete once TEER values were greater than 20Ω and showed stability over two readings 6 hours apart. Mature monolayers were treated with TNFα (20 ng / mL) or vehicle overnight immediately prior to final TEER measurements. All experiments were performed in triplicate, and empty (cell-free) filters were included in each run to measure baseline resistances.

### Transendothelial migration assays

HMVECs were plated on soft substrates and incubated as described above to allow formation of monolayers. Monolayers were monitored visually by phase-contrast imaging to ensure that they covered the substrate completely, without defects, before TEM assays were performed. Mature monolayers were treated with TNFα (20 ng / mL) overnight immediately prior to TEM assays. NK-92 MI cells were added to HMVEC monolayers in EGM-2 MV culture medium and placed into an environmental chamber (Stage Top Incubator, Tokai Hit, Shizuoka-ken, Japan) with 5% CO_2_ at 37ºC on an inverted microscope (Olympus IX83). DIC images were captured at 10-sec intervals with a 10X objective. Images of monolayers were analyzed using FIJI (Schindelin, 2012). Time-lapse movies were reviewed to follow individual tumor cells as they migrated into the endothelial monolayer.

## ACKNOWLEDGMENTS

This work was supported by NIH R35 grant GM144082 (J.A.C.). We are grateful to Joanna Kim and Olivia L Mooren for discussions and advice.

